# Comparative human embryo-mapping reveals neural bias of neuromesodermal progenitors in stem cell axial elongation models

**DOI:** 10.64898/2026.03.06.710064

**Authors:** Yuting Wang, Rafaella Buzatu, Cecile Herbermann, Micha Drukker, Christian Schröter

## Abstract

Elongation of the human body axis is driven by the coordinated differentiation of neural and mesodermal tissues. Pluripotent stem cell-based organoid models recapitulate key aspects of axial elongation, but how they relate to the regional organization of axial development in the embryo remains unclear. Here, we map scRNA-seq from twelve stem cell-based axial organoids to a stage-matched embryo. Integrated transcriptomic and trajectory analyses show that human axial development involves three origins: forebrain neural progenitors, primitive streak mesoderm, and neuromesodermal progenitors (NMPs) with posterior potency. Axial organoids capture subsets of these origins, and contain NMPs with strong neural bias. A multivariate regression model linking pathway modulation to cell type frequencies revealed a key role of TGF-β inhibition in the transition from anterior to posterior progenitors. Together, these findings clarify how the human body axis is partitioned into distinct domains by coordinated progenitor states and signaling.

## Introduction

The anterior-posterior (AP) body axis of vertebrates is built successively from head to tail. Despite its apparent continuity, the axis thus originates from continuously changing progenitors. Analyses in mouse and chick embryos have shown that the anterior neural tube which gives rise to forebrain, midbrain, and anterior hindbrain regions originates from anterior pluripotent epiblast ^1,2^. More posteriorly, epiblast cells that migrate through the primitive streak generate presomitic mesoderm (PSM) which subsequently forms somites that give rise to the cervical and upper thoracic vertebrae ^3–6^. Subsequently, the posterior neural tube and its adjacent somites, extending from the cervical spinal cord to the tail ^3,7^, arise from a distinct bipotent cell population located at the node–streak border (NSB), termed neuromesodermal progenitors (NMPs). After their initial specification, NMPs relocate progressively to more posterior stem-like niches, moving from the NSB at around embryonic day (E)7.5 to the caudal lateral epiblast (CLE) by E8.5. By approximately E10.5, NMPs become confined to the chordoneural hinge (CNH), where they terminally differentiate as axial elongation concludes around E13.5 ^8–10^. Little is known about the progenitor populations that underlie A-P axis development in human embryos, largely owing to limited access to post-implantation stages and their short-lived presence during early post-implantation development.

Cell allocation during the formation of the A-P axis in mouse embryos is orchestrated by combinations of extracellular signaling cues that promote distinct progenitor types. Anteriorly, antagonists of BMP and WNT pathways, including NOGGIN, CHORDIN, and DKK1, prevent mesodermal differentiation and instead allow direct neural differentiation of pluripotent epiblast cells ^11^. More posteriorly, BMP and WNT ligands, most notably BMP4 and WNT3, drive primitive streak formation and subsequent differentiation of the PSM that gives rise to cervical and upper thoracic vertebrae ^12–14^. Subsequently, in the NSB, high WNT and FGF signals from the primitive streak promote the specification and self-renewal of NMPs ^15^. The balance of WNT/FGF signaling within the NSB, as well as in the subsequent NMP niches, influences NMP differentiation. This occurs through modulation of *Sox2* and *T* (also known as *Brachyury* or *TBXT*) expression, transcription factors classically associated with neural and mesodermal fates, respectively ^6,16^. Mechanistically, high WNT3A/β-catenin signaling promotes *T* expression and PSM differentiation through positive feedback with FGF signaling, whereas reduced WNT3A favors *Sox2* expression and neural tube differentiation^10,16–18^. It remains unclear whether human NMPs exhibit comparable *SOX2/TBXT* heterogeneity and balance-driven differentiation.

A major advance for studying human axial development has been the establishment of protocols to generate axis-forming organoids from human induced and embryonic pluripotent stem cells (i/ePSCs). These human axial organoid protocols build on foundational mouse gastruloid studies demonstrating neural-tube and somite-like structures and progressive spatiotemporal *HOX* gene activation characteristic of axial elongation *in vivo* ^19–22^. By adjusting the timing, dosage, and duration of signaling regimes, protocols were tuned to promote specifically the development of neural tube (NT) ^23–25^, somite-derived mesodermal (SM) tissues ^24,26–28^, or their co-development in trunk-like (TK) structures ^24,29–32^. Despite differences in protocols and tissue outputs, a common feature of these axial organoids is that they contain NMP-like cells. However, the lineage potency of these NMPs as well as their contributions to somite and neural tube structures within the organoids has not been systematically analyzed across protocols.

Here, we address this question by systematically mapping scRNA-seq data of twelve human i/ePSC-derived axial organoid models, including time series datasets, to a human post-conception week 3 (PCW3) embryo reference that includes neural tube, PSM, somites, and NMP populations^23–30,32,33^. We combine transcriptomic alignment and RNA velocity–based trajectory inference to reconstruct differentiation trajectories across protocols. This analysis reveals that NMP-like populations in current human axial elongation protocols are predominantly biased toward neural identities and contribute minimally to posterior PSM-like cells. We also developed a mathematical model to dissect signaling pathway-level effects on tissue outputs from combinatorial signaling in protocols. This analysis reveals both known and novel signaling functions, including a critical role of TGF-β inhibition in the appearance of NMPs. Overall, our work suggests that current human axial organoids capture distinct submodules of a tripartite developmental program present in the embryo.

## Results

### Mapping single-cell transcriptomes from axial elongation models to a common human embryo reference

To analyze embryonic correspondences in axial organoids, we used scRNA-seq data from a PCW3 human embryo undergoing early axial elongation and containing SOX2⁺/TBXT⁺ NMPs as a reference (Fig. 1A) ^34^. To achieve fine-grained cell type resolution, we first re-clustered the reference dataset into 28 clusters that were annotated based on differentially expressed and marker genes in each cluster (Fig. 1B, Fig. S1, Table S1). These clusters contained the main subpopulations that contribute to head-to-tail axis formation, such as NMPs, forebrain–midbrain, hindbrain, and cervical–thoracic neural tube, as well as anterior and posterior PSM, intermediate mesoderm, lateral plate mesoderm, and mesoderm-derived muscle cells. Onto this reference, we projected scRNA-seq data from axial organoids generated with 12 distinct protocols, including time-series datasets (Fig. S2 and Table S2). Three protocols were reported to yield neural-tube tissue ^23–25^, four somite tissue ^24,26–28^, and five trunk-like structures containing both neural-tube and somite tissue ^24,29,30,32,33^.

**Figure 1.**
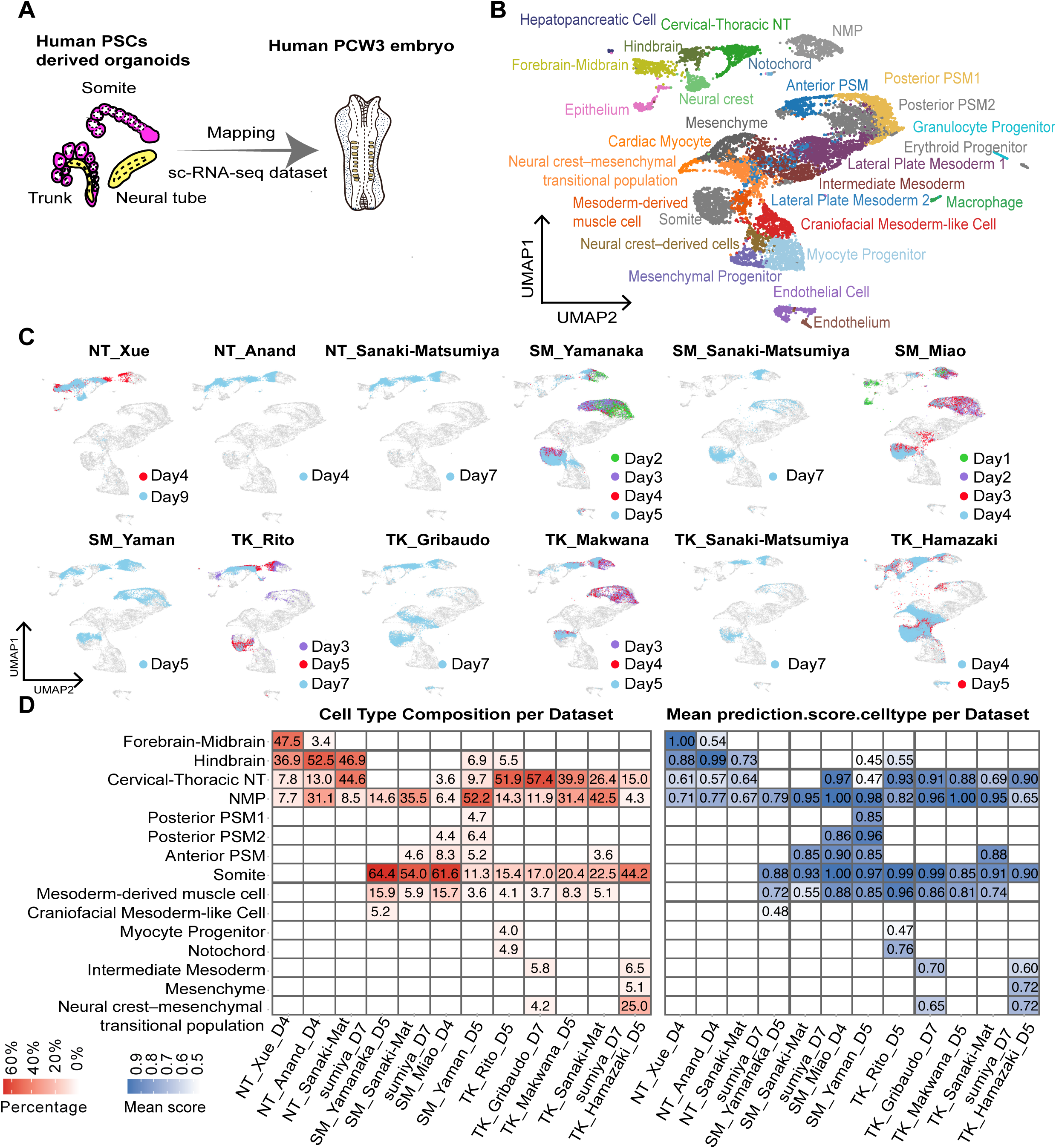
Mapping of organoid single-cell datasets with human PCW3 embryo. (A) Overview analysis strategy, showing the projection of existing human neural tube, trunk, and somite organoid single-cell datasets onto the PCW3 human embryo reference. (B) UMAP visualization of major cell populations from the PCW3 human embryo reference atlas. (C) Projection of each organoid dataset onto the PCW3 UMAP embedding. Time-points from time-resolved organoid datasets are color coded (D) Heatmap showing label transfer results. Left panel shows the percentage of predicted cell type labels obtained for each organoid, right panel displays the mean predicted score for each cell type. Cell populations representing fewer than 3% of the total cells were omitted for clarity.

For each protocol, we applied label transfer to assign embryonic identities to organoid-derived cells (Fig. 1C). We then quantified the percentage of assigned cell types and their associated confidence scores at day 5 of differentiation or the closest corresponding timepoint where scRNA-seq data was available (Fig. 1D). Across all protocols, confidence scores for the major populations were similarly high, supporting accurate assignment to the embryonic identities present during axial development (Fig. 1D). Notably, NMP-like populations were detected in all organoids with moderate to high confidence scores. Forebrain–midbrain like clusters were predominantly detected in NT_Anand, NT_Xue, SM_Miao, and TK_Rito organoids, with these clusters appearing only transiently in the latter two (Fig. 1D, Fig. S3B). Cells from NT_Anand, NT_Xue, and NT_Sanaki-Matsumiya organoids mapped primarily to hindbrain, cervical-thoracic neural tube and NMPs, with additional neural cells also detected in SM_Miao and SM_Yaman organoids. In contrast, neural cell types detected in TK protocols were almost exclusively classified as cervical-thoracic neural tube, indicating that these protocols promote more posterior neural identities compared with NT protocols. Fewer than 3% of the cells from the NT organoids mapped to mesoderm clusters.

In contrast, SM and TK organoids consistently generated somite cell populations and frequently exhibited mesoderm-derived muscle cells. Additionally, anterior and posterior PSM1/PSM2 populations, corresponding to immature mesodermal states, were present in some of the SM protocols but were only transiently observed in one of the five TK protocols. This indicates that the signaling cues in SM organoid protocols are more conducive to sustained generation of PSM cells, whereas TK organoid protocols appear to favor a single early wave characteristic of trunk somite formation. Consistent with this evidence for ongoing mesoderm differentiation, NMP confidence scores were higher in SM and TK than in NT organoids (Fig. 1D). Cells labeled as intermediate mesoderm were only detected in the TK_Gribaudo and TK_ Hamazaki datasets. While NMP-like cells were detected across all protocols, with proportions ranging from 5% to 50%, time-resolved datasets showed that the NMP abundance generally declined with differentiation time (Fig. S3B). This coincided with an increase in the abundance of cell types from more differentiated tissues such as hindbrain and cervical-thoracic neural tube in NT and TK protocols, and somite and muscle in TK and SM protocols (Fig. S3B). Collectively, these analyses show that axial organoid models incompletely and unevenly recapitulate the embryonic A-P axis, with individual protocols preferentially recapitulating to different axial levels and tissue types.

### NMPs in axial elongation models progressively acquire a neural bias

To investigate NMP bifurcation dynamics in human axial organoids, we next generated a focused reference projection of the PCW3 human embryo scRNA-seq data containing NMP-like cells together with their known embryonic progeny, including cervical-thoracic neural tube, anterior and posterior PSM1/2 and somites, the latter of which are also known derivatives of the primitive streak. We also included clusters annotated as forebrain-midbrain, hindbrain, and notochord to enable analysis of anterior neural development (Fig. 2A). As NMP fate decisions in mouse and chick are linked to a continuum of TBXT/SOX2 expression ratios ^35^, we examined whether a comparable gradient exists in the human embryo. Mapping TBXT and SOX2 expression in the PCW3 embryo onto the focused UMAP revealed a similar continuum of TBXT-to-SOX2 expression ratios across the NMP cluster (Fig. 2B). TBXT expression displayed a graded domain spanning NMPs and extending into posterior PSM1 and PSM2 clusters. Conversely, SOX2 was enriched in neural tube clusters and decreased toward the NMP region where TBXT levels were highest.

**Figure 2.**
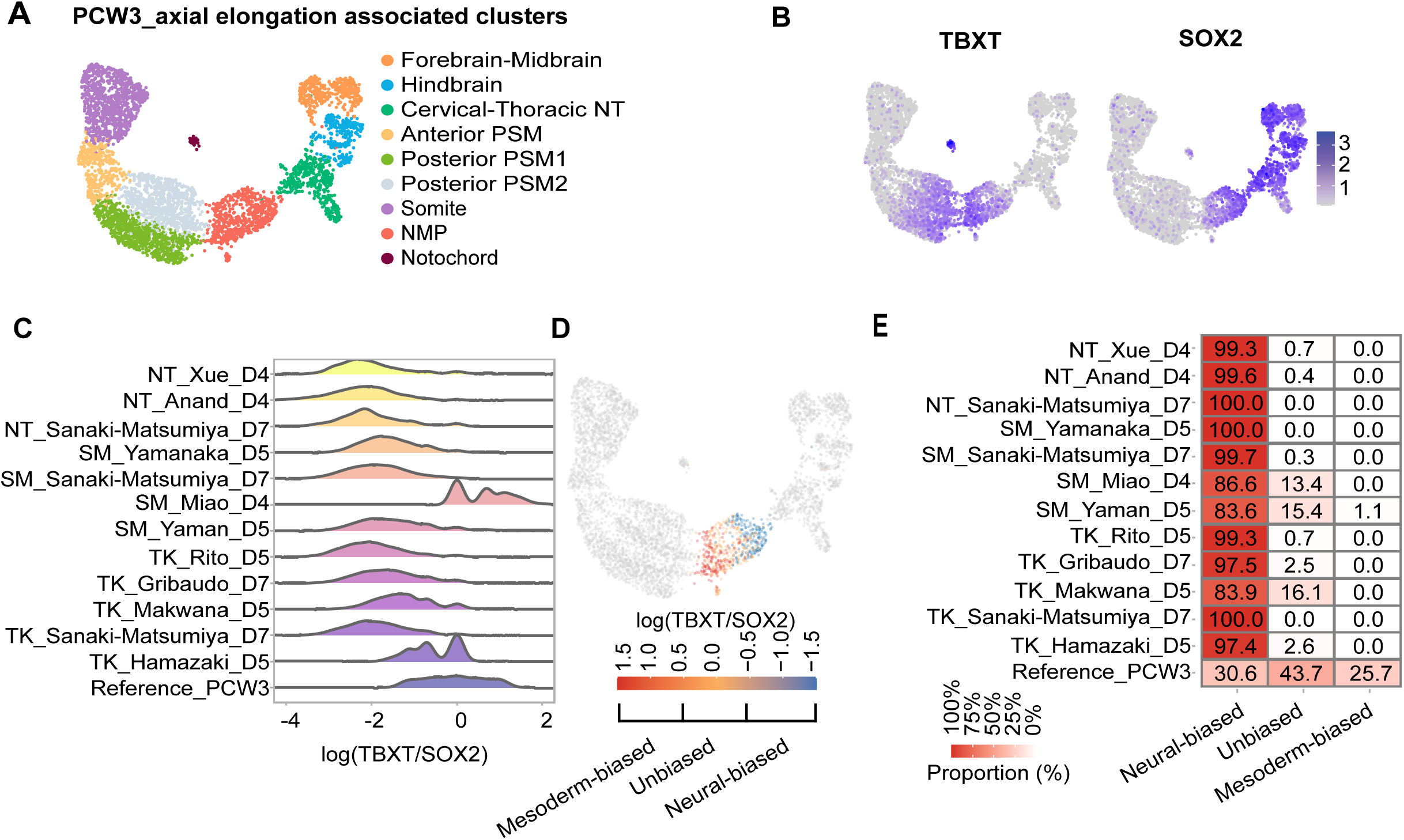
Neuromesodermal progenitors (NMPs) in axial elongation models acquire a neural bias. (A) Focused UMAP projection of selected cell populations from the human embryo reference, including NMP-derived populations (three PSM populations, somite, cervical-thoracic neural tube), forebrain-midbrain, hindbrain, and notochord cells. (B) Expression of TBXT and SOX2 across the selected populations. (C) Comparison of TBXT/SOX2 expression ratios between NMPs from the embryo and from various axial elongation models at selected timepoints. (D) Classification of embryonic NMPs into three subtypes based on TBXT/SOX2 expression ratio thresholds: putative unbiased (-0.5 ≤ ratio ≤ 0.5), mesoderm-biased (ratio > 0.5), and neural-biased (ratio < -0.5). (E) Proportion of each NMP subtype classified based on thresholds from (D) in axial elongation models and the embryo reference.

To determine how these transcriptional states are represented in axial organoids, we quantified TBXT/SOX2 expression ratios in NMPs (Fig. 2C). In nearly all organoids, NMPs exhibited lower TBXT/SOX2 ratios than embryonic NMPs, with the exception of SM_Miao, indicating a general neural bias in organoid-derived NMPs. Consistent with this, in time-series scRNA-seq datasets, we observed a progressive decline in TBXT/SOX2 ratios at later time points, accompanied by an increase in neural progenitors (Fig. S4A).

To quantify the distribution of NMPs across putative bipotent, mesoderm-biased and neural-biased states, we defined two TBXT/SOX2 expression thresholds separating the three classes (Fig. 2D). Applying the thresholds to the PCW3 human embryo revealed that 25.7% of NMPs were mesoderm-biased, whereas 43.7% and 30.6% were classified as unbiased and neural-biased, respectively (Fig. 2E). In contrast, mesoderm-biased NMPs were detected only transiently in organoid models, for example, between days 2-3 in SM_Yamanaka, days 1-3 in SM_Miao, and days 3-4 in TK_Makwana, and were largely depleted across protocols from day 4 onward (Fig. 2E, Fig. S4B). Among all organoids, only three of twelve, specifically, SM_Miao, SM_Yaman, and TK_Makwana, retained 10-20% unbiased NMPs at day 4 or later, whereas the remaining protocols displayed markedly smaller unbiased NMP populations. Conversely, neural-biased NMPs dominated across all organoid protocols, comprising more than 80% of the NMP population (Fig. 2E). This neural predominance further intensified over time, as longitudinal datasets revealed a progressive increase in neural-biased NMPs accompanied by a decline in bipotent and mesoderm-biased fractions (Fig. S4B). Together, these analyses indicate that NMPs in all axial elongation organoids progressively adopt a neural bias during differentiation, even when mesodermal derivatives dominate the final cell output.

### RNA velocity uncovers developmental trajectories shared between human embryo and axial organoids

Next, we asked whether organoid-derived NMPs progress toward the same fates, and along matching developmental trajectories, as embryonic NMPs. Using eleven organoid-derived datasets with unspliced–spliced transcript data, we computed RNA velocity vectors and re-embedded cells onto the axial-focused PCW3 embryo UMAP with transferred cell type identity labels. In the PCW3 embryo, RNA velocity vectors revealed two opposing streams from the NMP cluster, one directed toward the cervical-thoracic neural tube and another toward the posterior PSM populations (Fig. 3A). While the neural trajectory extended into part of the hindbrain cluster, remaining velocity vectors within the hindbrain and forebrain-midbrain clusters appeared discontinuous and spatially segregated from the NMP-derived stream. This pattern is consistent with anterior pluripotent epiblast progenitors giving rise to midbrain-forebrain and anterior hindbrain domains, while NMPs contribute to neural tube formation from the posterior hindbrain through the tailbud ^6^. The mesoderm stream is also consistent with anterior and posterior PSM populations that progressively mature into somites ^3–5^, as reflected by velocity vectors linking the anterior and posterior PSM1/PSM2 clusters to the somite cluster. However, only sparse vectors extended from the NMP cluster toward this mesodermal stream, indicating that the contribution of NMPs to paraxial mesoderm is limited at PCW3. Interestingly, a third set of velocity vectors was confined to the NMP cluster, plausibly reflecting self-renewal within the NMP compartment.

**Figure 3.**
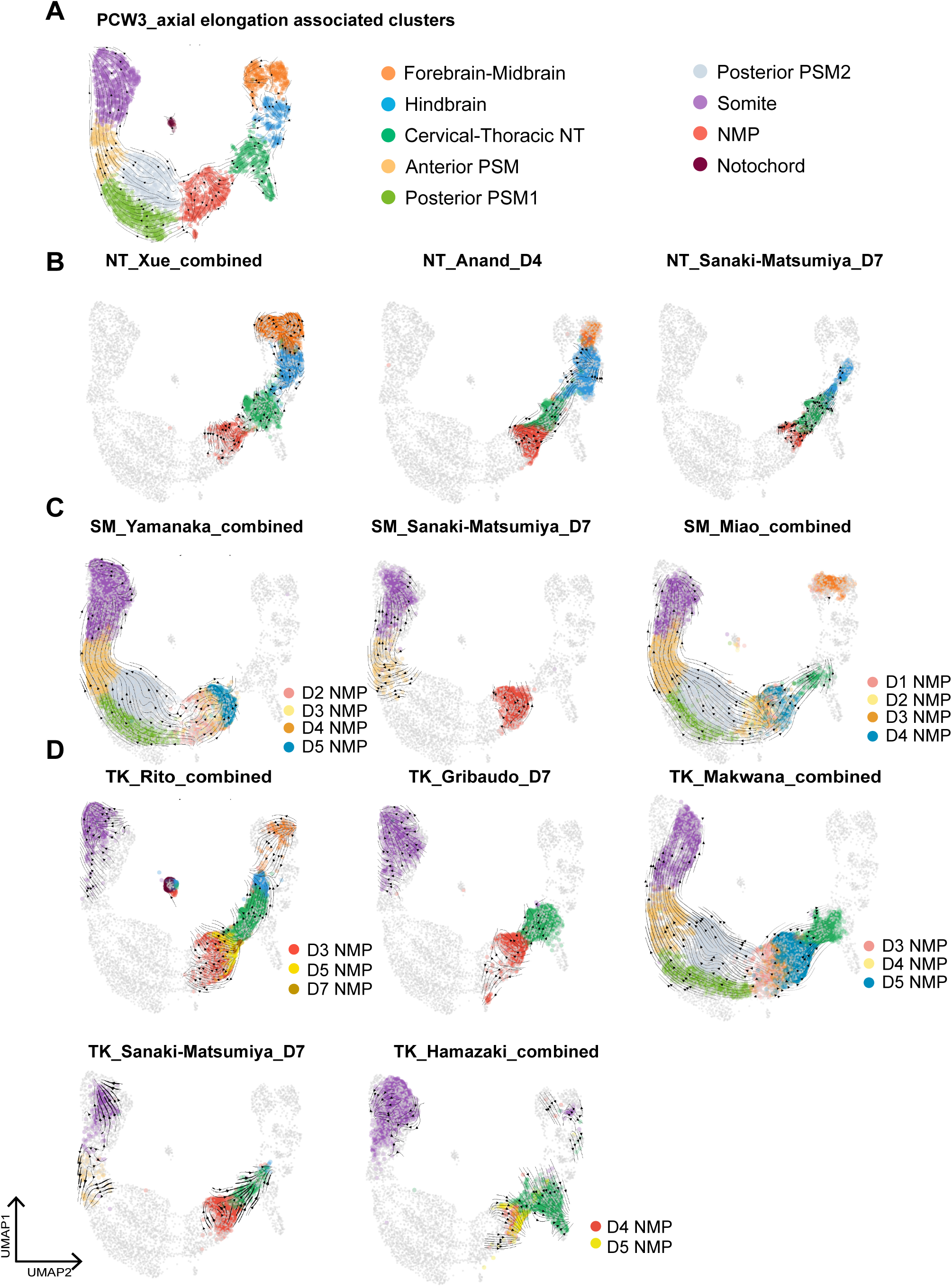
Contribution of NMPs to neural and somitic tissues in human embryos and axial organoids. (A) Focused UMAP with RNA velocity vectors of embryo reference. (B) Axis-associated cell types and RNA velocity vectors of neural tube (NT) organoids projected onto focused embryo reference. (C), (D) Same as (B) but for somite organoids (C) and trunk-like structures (D).

Inspection of velocity vectors in NT organoids revealed an embryo-like forebrain-midbrain to hindbrain stream and an NMP-to-cervical-thoracic neural tube stream (Fig. 3B). However, the relative prominence of these streams differed across NT protocols. NT_Xue displayed both streams, with a stronger forebrain–midbrain to hindbrain trajectory. NT_Sanaki-Matsumiya predominantly exhibited the NMP-to–cervical–thoracic neural tube trajectory, whereas NT_Anand showed contributions from both. These patterns indicate that the NT protocols capture largely overlapping stages of neural tube development, with NT_Sanaki-Matsumiya exhibiting most restricted cervical-thoracic neural tube differentiation.

The embryonic continuum of RNA velocity vectors extending from anterior and posterior PSM into somites was recapitulated in two somite models, SM_Yamanaka and SM_Miao (Fig. 3C). In contrast, SM_Sanaki-Matsumiya lacked immature PSM clusters and instead displayed a truncated velocity pattern, consistent with a more advanced somitic stage. The connection between NMPs and posterior PSM clusters was largely confined to early differentiation from approximately days 2-4. From day 4 onward, RNA velocity vectors in NMPs from SM_Sanaki-Matsumiya and SM_Miao organoids predominantly pointed toward, or remained within, the neural trajectory, further supporting the conclusion that even in SM protocols, NMPs progressively acquire a neural bias.

Connections from NMPs to PSM clusters were also limited in the TK models, as four of the five organoids generated minimal or no PSM. Only TK_Makwana showed trajectories from NMPs toward PSM around day 3 (Fig. 3D). In contrast, vectors extending from NMPs toward neural clusters were consistently detected across TK organoids, although generally less prominently than in NT organoids. Because projecting RNA velocity vectors onto a UMAP generated from another dataset can introduce distortions and potentially lead to spurious alignment with the reference structure (“force-mapping”), we also computed RNA velocity within each organoid dataset’s native embedding (Fig. S5). These analyses recapitulated the same trajectory structures, supporting the validity of the projection-based results. Taken together, the results indicate that human PSC-derived axial organoids recapitulate selected developmental trajectories observed in the human embryo. NMPs consistently contribute to posterior neural tube formation, whereas their contribution to somitogenesis appears temporally restricted, likely reflecting to the transient presence of TBXT^high^ NMPs in these organoids.

### Multivariate modelling reveals signaling pathway-specific influences on axial domain specification

Finally, we developed a quantitative model linking signaling modulation to A-P axis regionalization. To this end, we categorized hindbrain and cervical–thoracic neural tube populations as posterior neural tube (NT) tissue, forebrain-midbrain populations as anterior NT tissue, posterior PSM1/PSM2, anterior PSM, somites, myocyte progenitors, mesoderm-derived muscle cells, and craniofacial mesoderm-like cells as mesodermal tissue, and considered NMPs as a distinct tissue domain. Principle component analysis (PCA) of the twelve protocols based on the frequencies of the four tissue domains segregated the organoids in a manner consistent with the primary tissue types reported in the original publications (Fig. 4A). Next, we computed modulation scores for the five signaling pathways used across protocols, incorporating both concentration and exposure duration (Fig. 4B, Materials and methods). For signaling pathways targeted by multiple reagents, namely FGF, SMAD and RA, we computed pathway scores by summing scaled agonist and antagonist contributions (Fig. 4B). Moreover, inhibitors of BMP and Nodal-signaling were combined into a single SMADi score. PCA of signaling modulation scores showed that the twelve protocols did not segregate according to their primary tissue types (Fig. 4C). This was reflected in PC loadings, as both PC1 and PC2 were influenced by multiple signaling pathways. PC1 was driven mainly by FGF, RA, SMADi and SHH, whereas PC2 was dominated by SMADi, WNT, and RA contributions (Fig. 4C). Consistently, plotting the frequencies of cells assigned to the four embryonic tissues relative to pathway activity scores revealed no robust correlations (Fig. S6). These findings indicate that differences in lineage output are not driven by the activity of individual pathways, but arise from coordinated changes across multiple signaling pathways.

**Figure 4.**
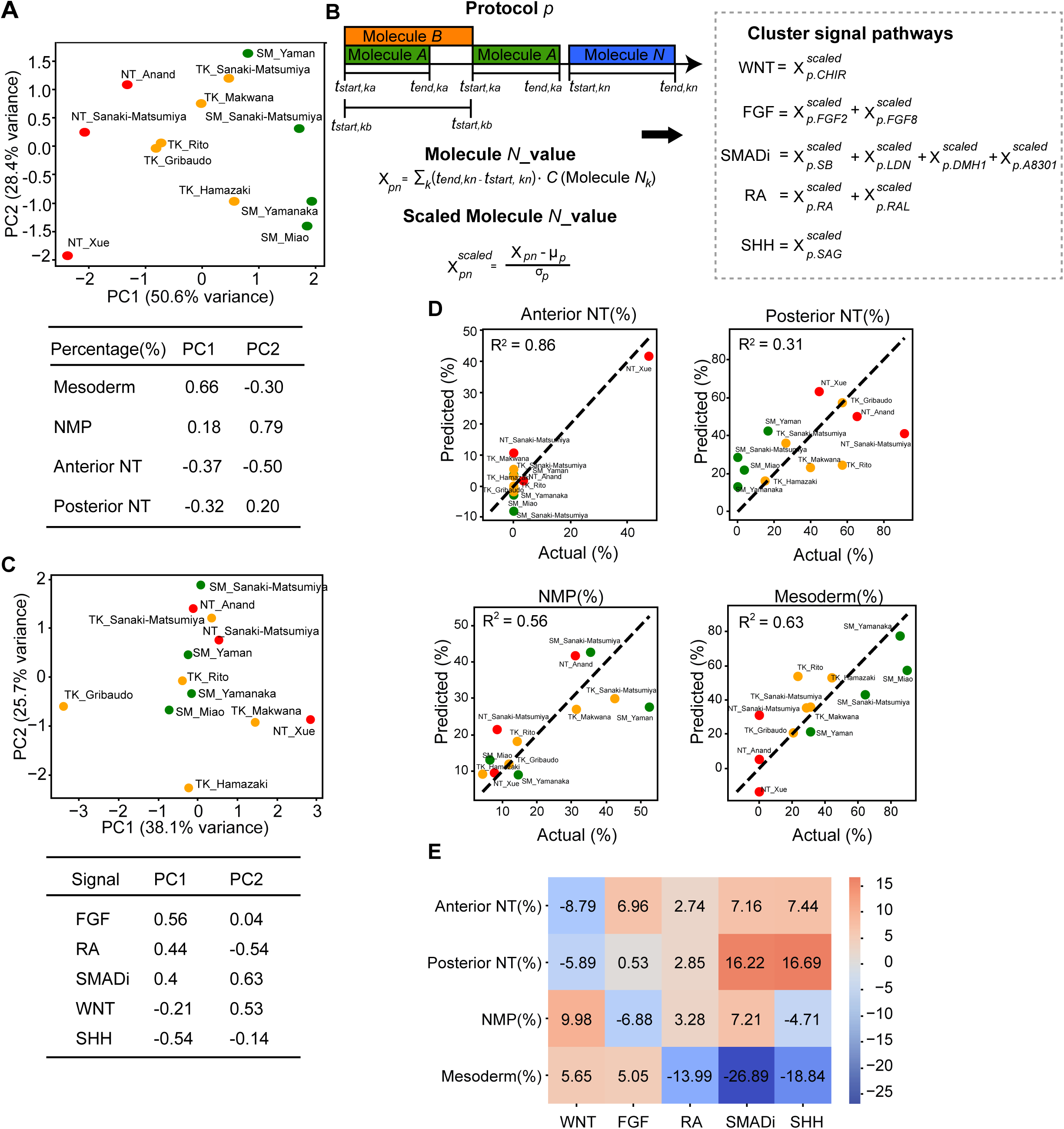
Multivariate modeling of combinatorial signaling contributions to lineage specification. (A) Principal component analysis (PCA) of twelve axial organoid protocols based on the proportions of four main tissue types - posterior NT, anterior NT, mesoderm, and NMPs. Loadings for the first two principal components are shown below. (B) Scheme of standardized pathway-level signaling score calculation. For each small molecule or growth factor treatment, the product of concentration and exposure duration was computed and then scaled. Scaled values were subsequently aggregated into pathway-level scores as follows: WNT = scaled CHIR; FGF = scaled FGF2 + scaled FGF8; SMADi = scaled SB + scaled LDN + scaled DMH1 + scaled A8301; RA = scaled RA + scaled RAL; and SHH = scaled SAG. (C) Principal component analysis (PCA) of standardized pathway-level signaling features across differentiation protocols and the corresponding first two PC loadings. (D) Goodness-of-fit of multivariate models for each tissue type outcome, quantified by coefficient of determination (R²) and mean absolute error (MAE). (E) Heatmap of multivariate linear regression coefficients for each tissue type outcome (anterior NT, posterior NT, NMP, mesoderm) and individual signaling pathway.

To quantify the relative contribution of each signaling pathway to the frequency of the cells assigned to the four embryonic tissues we used multivariate linear regression. This approach models cell frequencies per axis domain as weighted combinations of multiple signaling inputs, with weights estimated by parameter fitting. Across all four tissue categories, predicted frequencies showed moderate to strong agreement with observed cell frequencies (R^2^ between 0.31 and 0.86, Fig. 4D). Because a small number of protocols yielded anterior NT, the correlation is less informative for this tissue than for the other three, where predicted and observed frequencies are more broadly dispersed. The model’s ability to predict tissue frequencies indicates that combinatorial signaling inputs account for a substantial fraction of the variance in A-P axis regionalization.

We next examined the associations between signaling pathway weights, reflecting either positive (activating) or negative (repressive) contributions, and tissue outputs (Fig. 4E). Importantly, no single pathway uniformly explained domain allocation, reinforcing the combinatorial nature of signaling control. Activated WNT/β-catenin signaling showed a positive association with the presence of NMPs and mesoderm differentiation, but a strong negative association with both anterior and posterior NT tissues. Dual SMAD inhibition (SMADi) showed a positive association with anterior NT tissue, reinforcing the established role of BMP and TGF-β inhibition in anterior neural development ^6^. Notably, SMADi and RA were also positively associated with NMP frequency, with the RA association potentially reflecting the neural-biased state of NMPs in axial organoids. Unexpectedly, FGF signaling was negatively associated with NMP frequency, yet showed a clear positive association with mesoderm differentiation. These findings suggest that, within axial organoids, FGF primarily promotes the transition of NMPs toward mesodermal fates, thereby reducing their relative abundance. Moreover, the regression analysis highlighted TGF-β pathway inhibition as a contributor to NMP and posterior NT tissue abundance. Together, these results demonstrate that axial regionalization across differentiation protocols is shaped by combinatorial signaling inputs, with distinct and quantifiable contributions from individual pathways.

## Discussion

The growing sophistication of human axial organoid models provides an unprecedented opportunity to dissect the logic of human axial development that we exploit through the systematic and harmonized analysis of single cell data from multiple i/ePSC-derived axial organoid systems. By integrating scRNA-seq data with a stage-matched human embryo reference, we reveal organizational principles. In addition, our harmonized cross-protocol analysis enables a quantitative framework that explains how distinct signaling configurations regulate anterior–posterior tissue output. This represents one of the first systematic modeling efforts that quantitatively links signaling modulation to organoid outcomes.

Mapping results, differentiation trajectories inferred from RNA velocity analysis, and multivariate modeling collectively point toward a modular framework for human axial development (Fig. 5). We propose that discrete modules of embryonic axial development along the anterior–posterior (A–P) axis are represented to varying extents by different axial organoid systems. The anteriormost module underlies the formation of anterior neural tissues of the forebrain-midbrain and portions of the hindbrain. This module is most prominent in NT protocols, NT_Xue and NT_Anand, but also detectable in TK_Rito and in SM_Miao. More posteriorly, a mesodermal module generates the majority of mesodermal cells observed in SM and TK protocols. Because these cell types lack clear lineage connections to the NMP population, they likely represent anterior somites that arise from primitive streak mesodermal progenitors ^3–5^.

**Figure 5.**
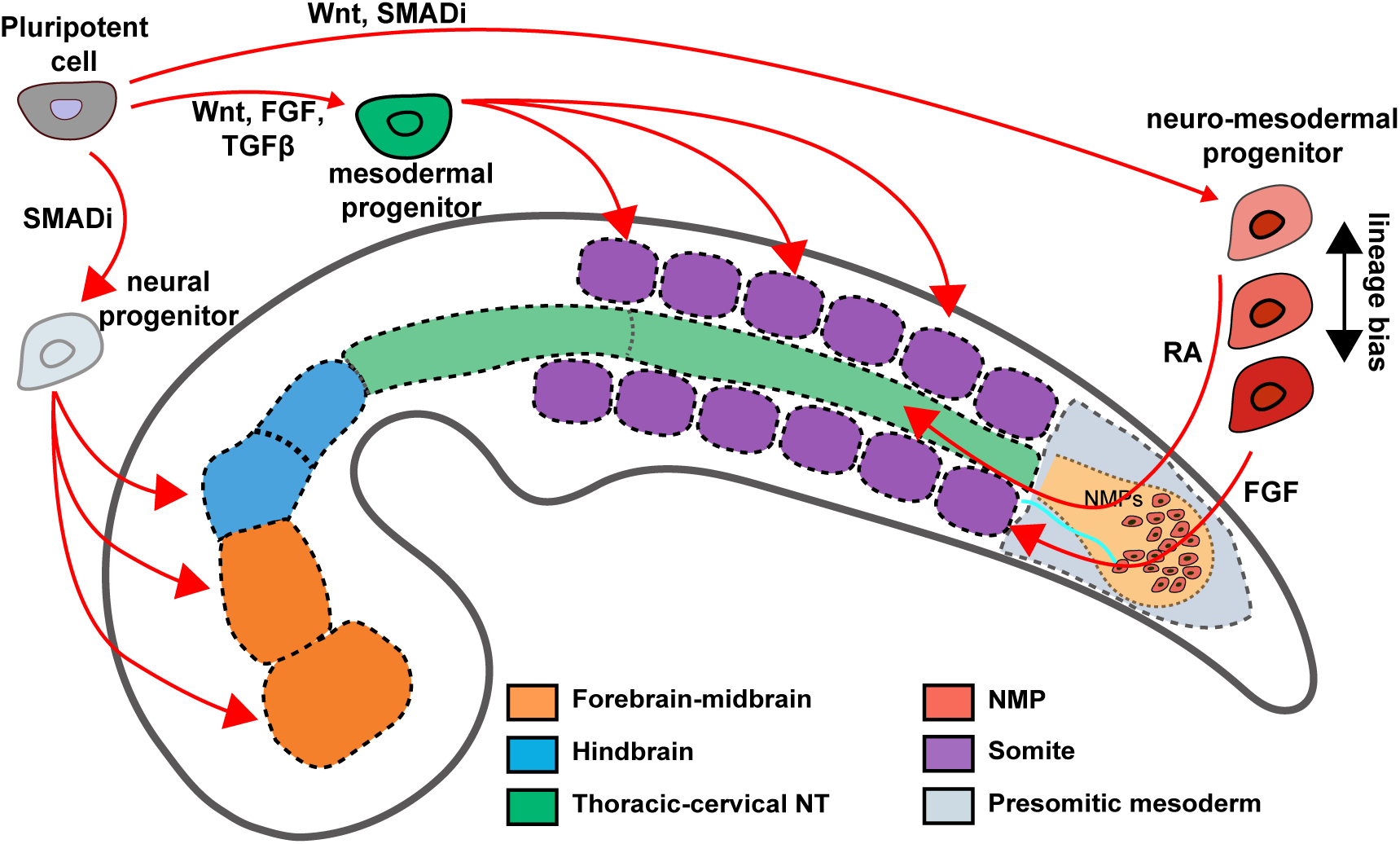
Modular framework for human axial development. Scheme indicates the progression of pluripotent cells to distinct types of progenitors and their tissue descendants along the developing axis. Associated signaling pathways are indicated next to differentiation trajectories.

The most posterior module is centered on NMPs and underlies the formation of posterior neural tissue from the hindbrain caudally, as well as more posterior somites. Notably, in both the embryo reference and in axial organoids, the contribution of NMPs to somitic tissue appears limited. This suggests that, at the PCW3 stage, NMPs predominantly generate neural output, some of which may be flanked by primitive streak-derived mesoderm. While embryonic NMPs likely acquire greater competence to generate posterior somites at later stages, efficient bi-lineage contribution is not observed in any of the axial organoids analyzed here.

A modular framework of human axial development is concordant with observations in chick and mouse embryos, where anterior somites arise from primitive streak–derived mesoderm, whereas NMPs primarily derive posterior neural tube extension and contribute to paraxial mesoderm during posterior somitogenesis. Thus, modular organization of axial development appears to be a conserved feature of amniote embryos, despite pronounced interspecies differences in early embryonic size and morphology. A key feature of modular axis development is that both neural and mesodermal cells, despite converging identities and a continuous arrangement along the axis, are specified through multiple routes that each rely on separate progenitors. Extending this framework to human development continues the challenge of the classical germ layer theory of gastrulation that began with the identification of NMPs in the mouse more than 15 years ago ^36^.

Finally, we introduced a quantitative framework that converts time and dose dependent signaling inputs across studies into standardized pathway-level features, and integrates them with unified quantifications of tissue frequencies derived from reference mapping. This integration enables systematic identification of signaling contributions to the three anterior–posterior developmental modules. The model recapitulates several well-established relationships from 2D differentiation systems, including the dependence of anterior neural differentiation on sustained SMAD inhibition ^37^, and the promotion of mesoderm differentiation by WNT and FGF signaling ^38^. Although our pathway-level quantifications do not capture detailed temporal dynamics of signaling exposure, and the current number of datasets is likely too limited for fully constraining all model parameters, the recovery of known signaling relationships substantiate the validity of the multivariate modelling strategy.

Importantly, our analysis reveals an additional role of SMAD inhibition in axial module allocation. While SMAD inhibition exerts a strong negative influence on the predominantly anterior mesodermal populations observed in axial organoids, it is positively associated with NMP frequency and posterior NT. This insight is consistent with recent findings from mouse gastruloids describing an anterior Nodal-dependent module and a posterior, Nodal-independent module ^39^. Notably, while we find that human axial elongation organoids preferentially generate anterior tissues, gastruloids made from mouse embryonic stem cells readily generate posterior trunk and tail tissues without TGF-β inhibition ^19^. We speculate that this important species-specific difference is caused by higher paracrine TGF-β signaling in human compared to mouse pluripotent cells. Such high baseline TGF-β signaling in human cells aligns with the use or the TGF-β pathway inhibitor SB431542 for generating self-renewing human axial stem cells that represent posterior identities ^40^.

More broadly, our approach for identifying quantitative relationships between signaling modulation and cell-type frequencies may serve as a general strategy to integrate data across diverse published i/ePSC differentiation protocols for elucidating mechanisms of human development.

## Supporting information

Table S1. Differentially expressed genes (DEGs) of human PCW3 scRNA-seq atlas

Table S2. Composition of differentiation media used across organoid protocols

Table S3. List of datasets included in this study

## Supplementary Figures

**Supplementary Figure 1.**
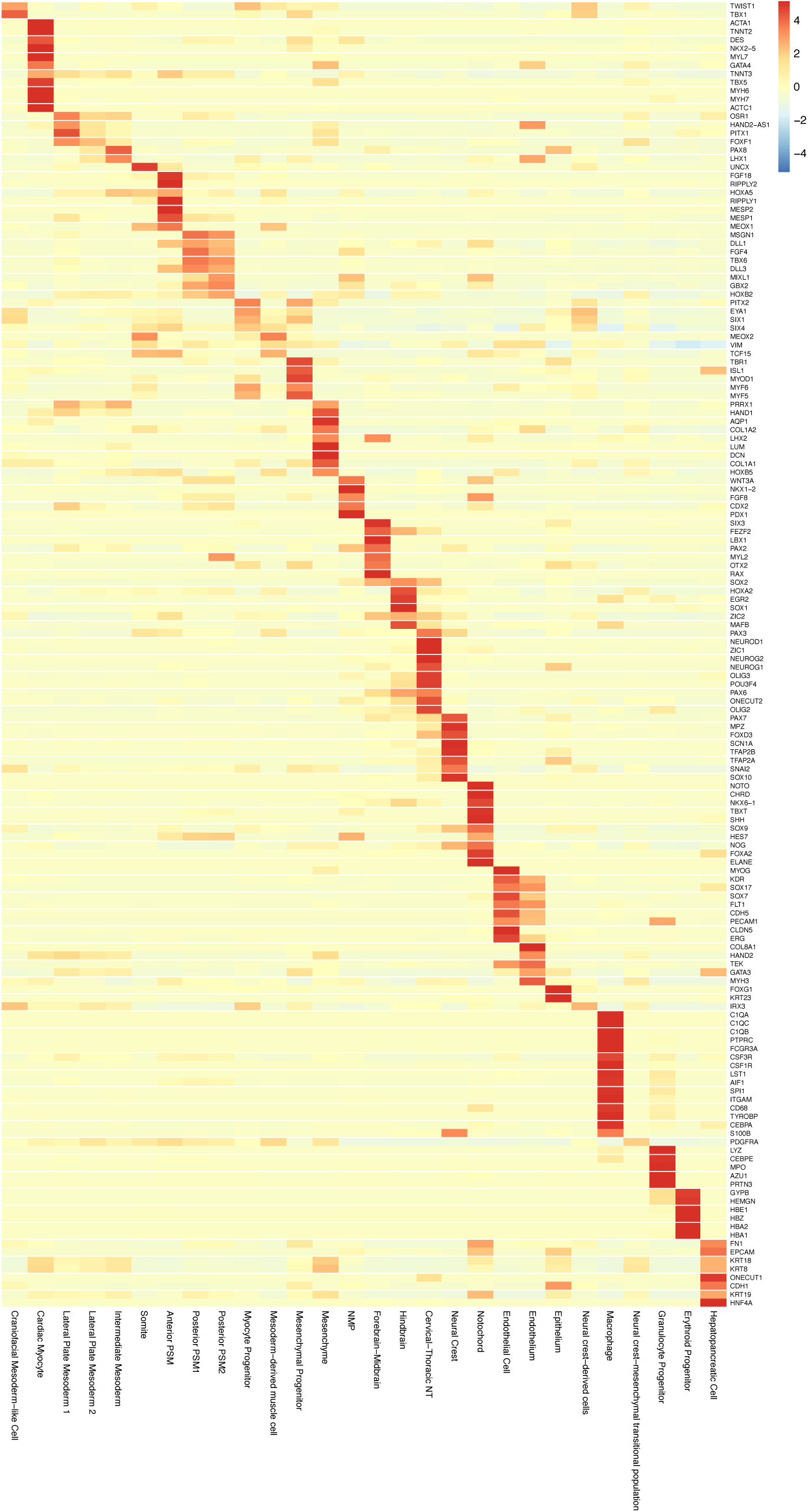
Heatmap showing expression of selected marker genes used to re-annotate cell types in human embryo reference.

**Supplementary Figure 2:**
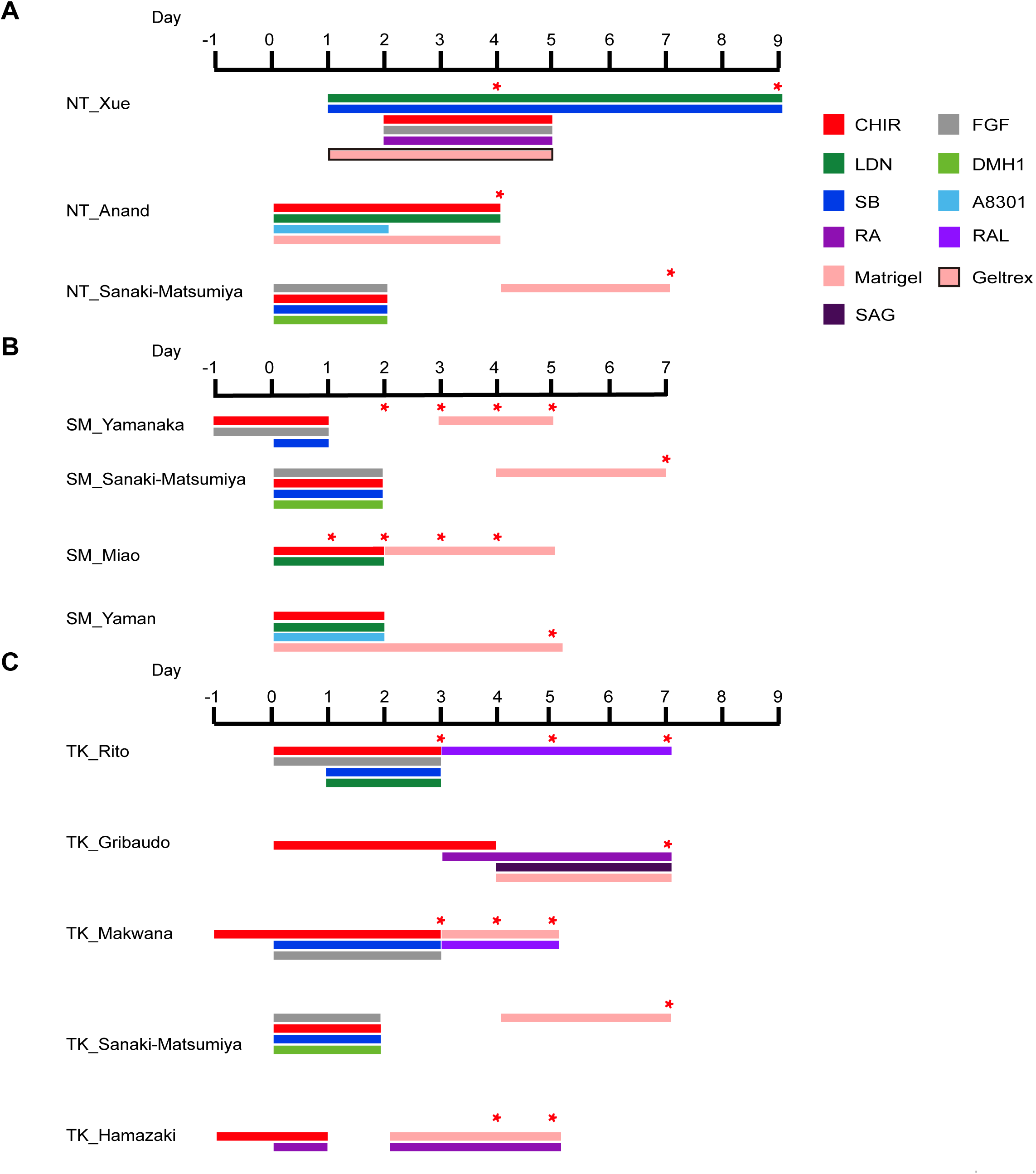
Graphical overview of hPSC-derived axial organoid protocols. Protocols are grouped by the predominant tissue type generated: neural tube (NT, panel A), somitic tissue (SM, panel B), or trunk-like structures containing both neural and somitic tissue (TK, panel C). Signaling modulations for each protocol are represented by grouped horizontal bars, wherein each colored segment indicates the presence and duration of specific small molecules or supplements applied during the differentiation process; CHIR (red), FGF (gray), LDN (dark green), DMH1 (light green), SB (dark blue), A8301 (light blue), RA (dark purple), RAL (light purple), Matrigel (pink), Geltrex (pink with black border), and SAG (purple-black). For specific concentrations of each compound see Table S2. Asterisks indicate the time points at which single-cell sequencing data were collected for each protocol.

**Supplementary Figure 3.**
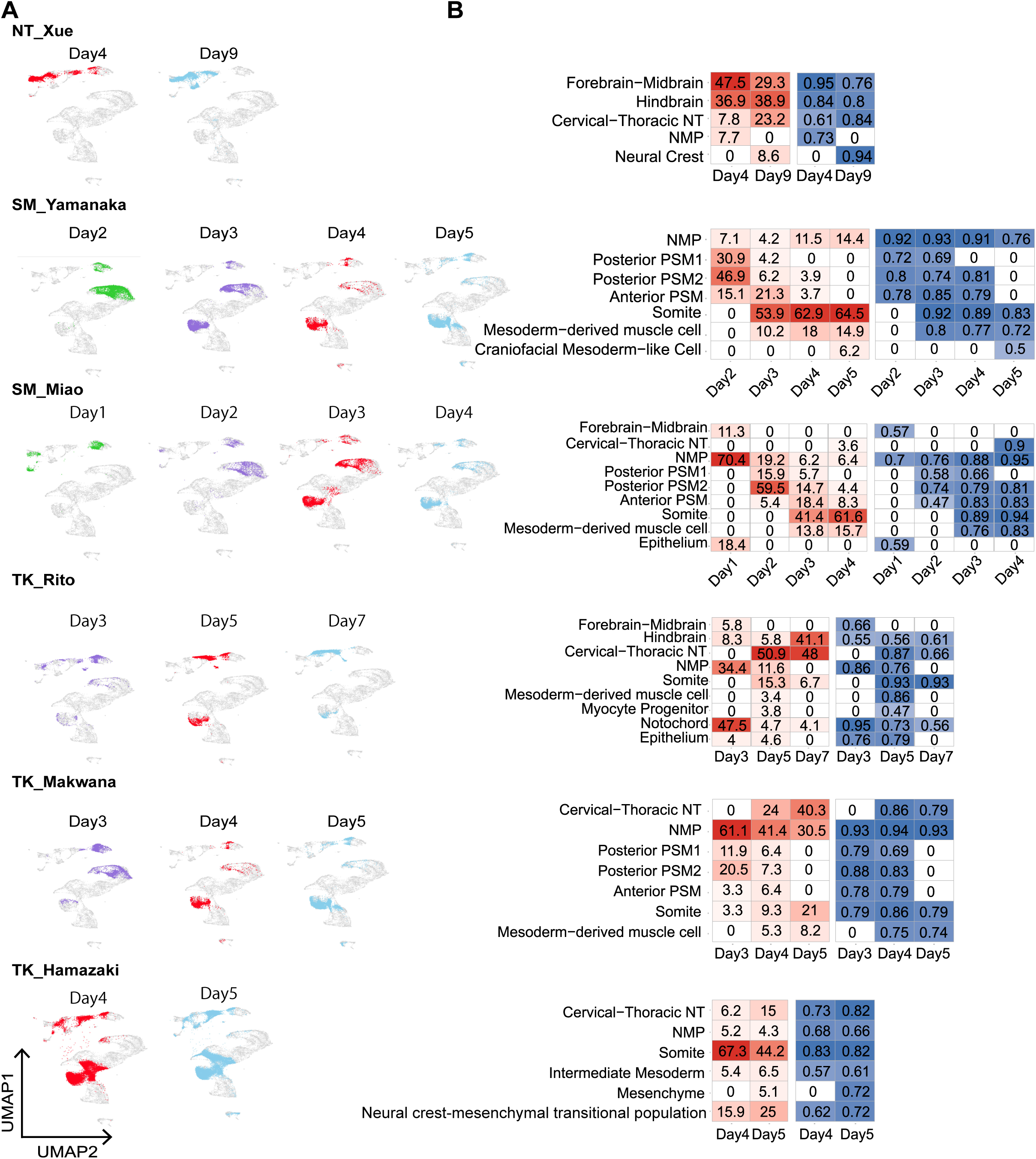
Time-resolved mapping of organoid single-cell datasets with human PCW3 embryo. (A) Time-resolved projection of single cell sequencing data from individual axial organoids onto the PCW3 UMAP embedding. (B) Heatmap summarizing the label transfer results based on transcriptomic similarity. Left panels show the percentage of predicted cell type labels obtained for each organoid at each timepoint (in red); right panels display the mean predicted score for each corresponding cell type (in blue). Cell populations representing fewer than 3% of the total cells were omitted for clarity.

**Supplementary Figure 4.**
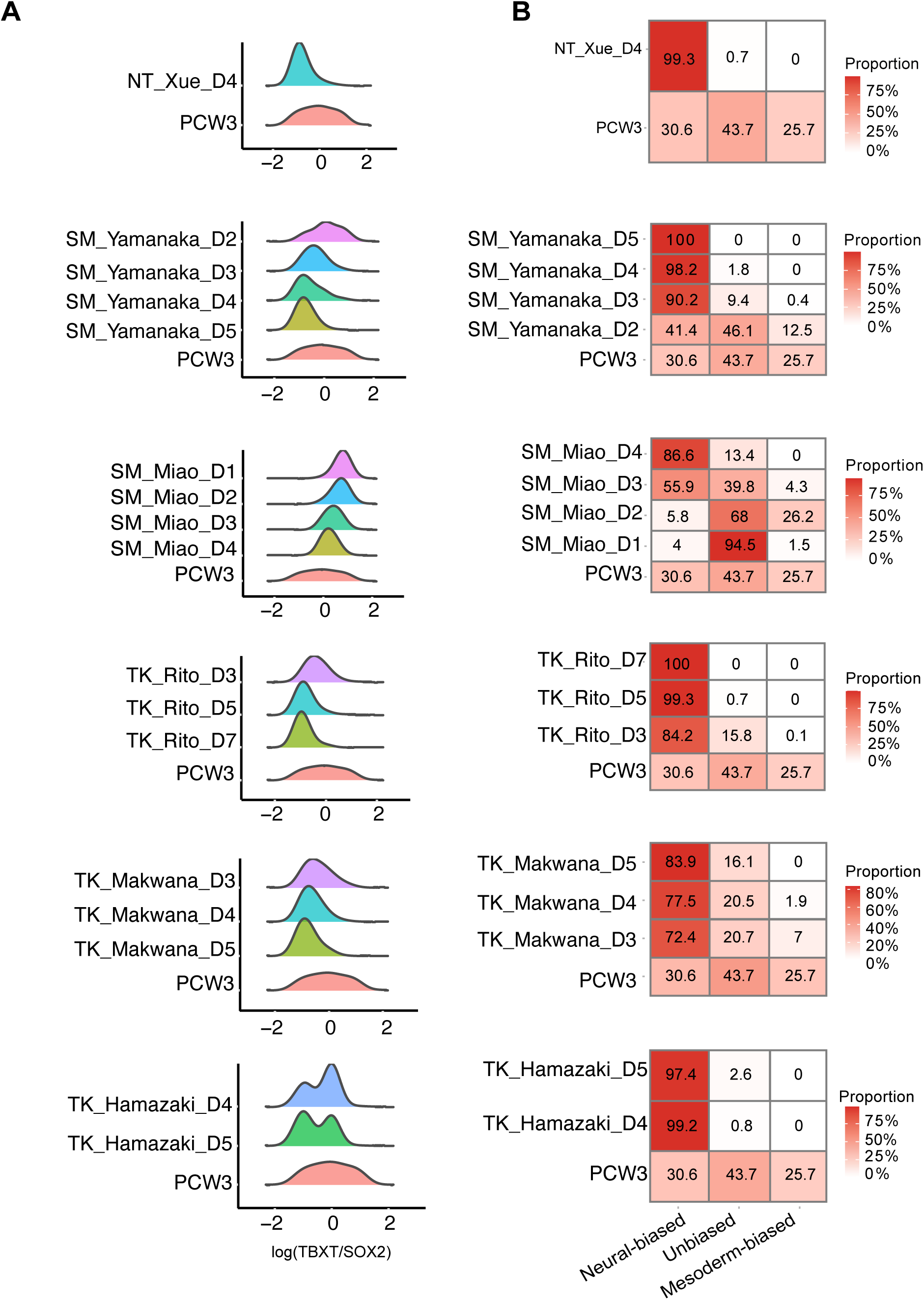
Time-resolved distribution of TBXT/SOX2 expression ratio in NMPs in individual organoids. (A) Ridgeplot showing of TBXT/SOX2 expression ratios in NMPs from the embryo and from various axial organoids at indicated timepoints. (B) Heatmaps showing proportions of putative bipotent NMPs, mesoderm-biased progenitors, and neural-biased progenitors as determined by TBXT/SOX2 expression thresholds shown in Fig. 2C for indicated timepoints and the embryo reference.

**Supplementary Figure 5.**
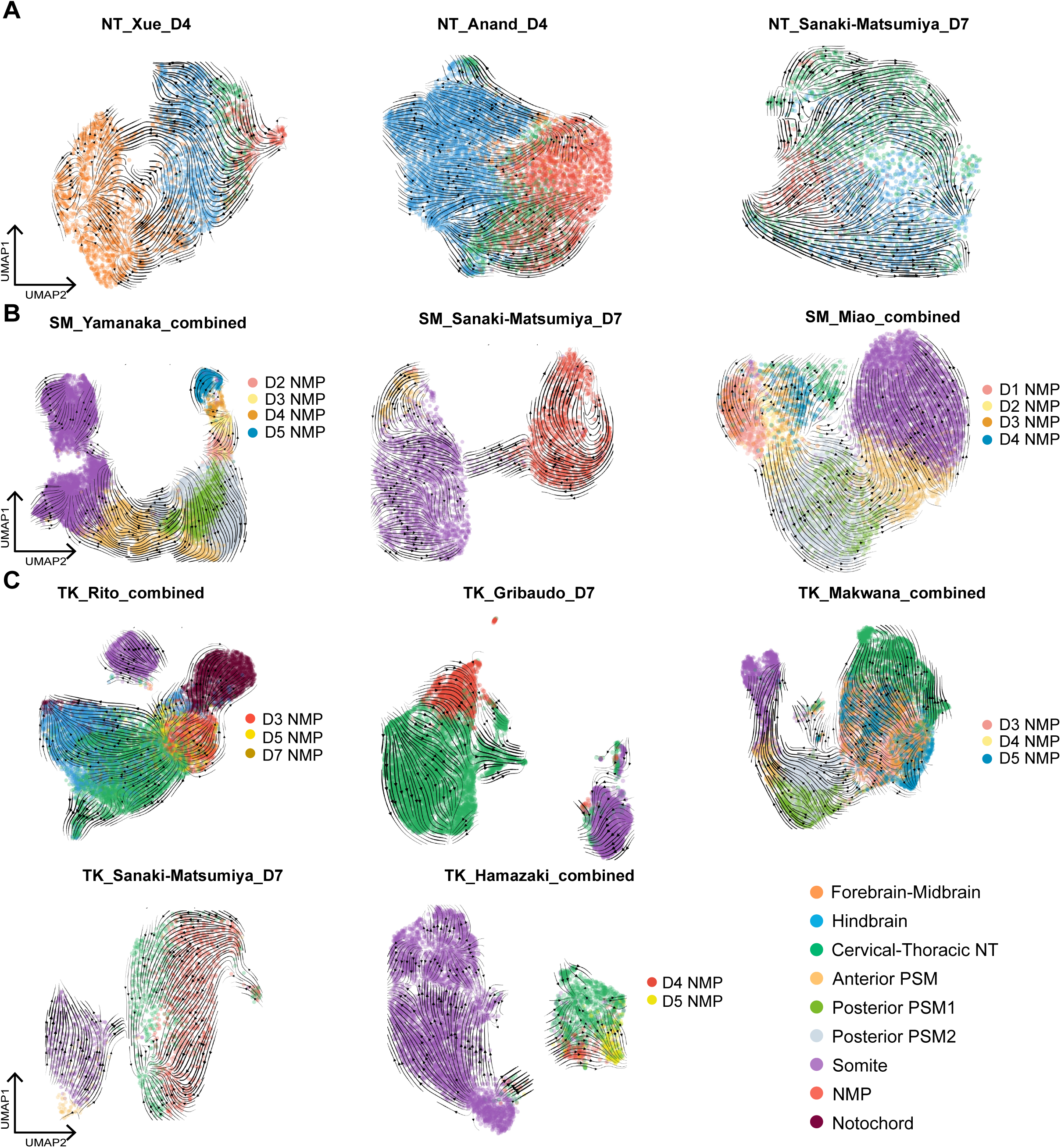
Individual UMAP projections and RNA velocity vectors of axial tissues from axial in organoids. Separate UMAPs for each axial organoid where spliced-unspliced transcript information was available each generated from the same cell types as in Fig. 3 (three PSM populations, somite, cervical-thoracic neural tube, forebrain-midbrain, hindbrain, and notochord). NMP populations are color-coded according to analysis time point; for all other cells, colors indicate cell type label obtained from label transfer. UMAPs are grouped according to main tissue type present: Neural tube (NT) organoids (A); Somitic (SM) organoids (B); and Trunk-like (TK) organoid (C).

**Supplementary Figure 6.**
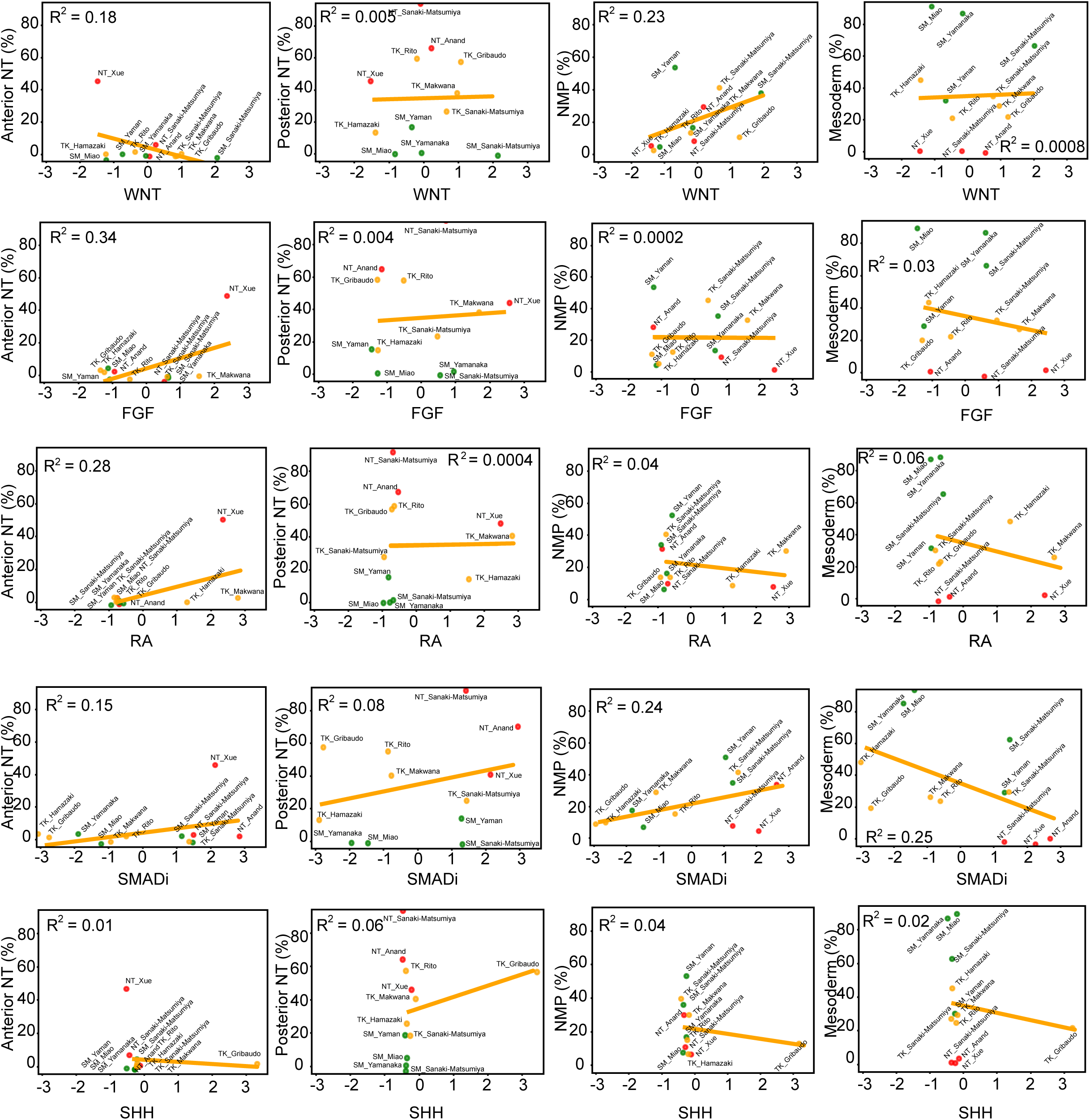
Univariate linear regression analysis of individual signaling pathways across four lineage categories. No significant correlations were detected for any single pathway in any of the lineage categories.

Table S1. Differentially expressed genes (DEGs) of human PCW3 scRNA-seq atlas

Table S2. Composition of differentiation media used across organoid protocols

Table S3. List of datasets included in this study

## Methods

### Datasets curation and preprocessing of the included scRNA-seq data of organoid

We included 12 human axial elongation organoid datasets from 9 studies^23–27,29,30,32,33^. The datasets analyzed in this study, along with their corresponding accession numbers, are listed in Table S3. For the NT_Sanaki-Matsumiya and TK_Sanaki-Matsumiya datasets, demultiplexing was performed following the procedures described in the original publication prior to further processing^41^. Raw FASTQ files of organoid datasets were processed using Cell Ranger (v9.0.1, 10x Genomics). Reads were aligned to the human reference genome (GRCh38), and the cellranger count pipeline performed read alignment, UMI quantification, and barcode filtering to generate gene-by-cell expression matrices. Aligned BAM files were retained for subsequent RNA velocity analysis, where spliced and unspliced transcript information was extracted to build loom files for downstream velocity inference. Each dataset was filtered using custom thresholds for gene and UMI counts and mitochondrial gene content (Table S3). Ribosomal genes were excluded from all datasets to reduce technical bias. Transcript counts from all datasets were normalized using Seurat’s LogNormalize method with a scale factor of 10,000 and natural log-transformed. The 2,000 most highly variable genes were selected with FindVariableFeatures, and the data were scaled using ScaleData with default parameters. Principal component analysis (PCA) was performed on the scaled data using the default 50 components.

### Clustering and annotation of the human PCW3 atlas scRNA-seq

Good quality cells were kept based on criteria listed in Table S3. Next, the filtered gene expression matrix was normalized and log-transformed. Highly variable features were identified using the “vst” method, selecting the top 2,000 variable genes. Data were then scaled, and principal component analysis (PCA) was performed using the top 2,000 variable features. For downstream clustering, the first 30 principal components were used to construct a shared nearest neighbor (SNN) graph with Seurat’s FindNeighbors, and clusters were identified using the FindClusters function with a resolution of 1. A total of 28 clusters were obtained. These were annotated based on the top 20 most differentially expressed genes in each cluster as well as the expression of known marker genes (Table S1, Fig. S1). Dimensionality reduction was performed using Uniform Manifold Approximation and Projection (UMAP) based on the first 30 principal components (RunUMAP) with the “umap-learn” method, and the UMAP model was retained for visualization.

### Query axial elongation organoid cells label transfer to PCW3 atlas

Cell type annotations from the PCW3 reference dataset were transferred to the query dataset using Seurat’s anchor-based integration framework ^42^. Briefly, anchors were identified as mutual nearest neighbors (MNNs) between reference and query cells based on k-nearest neighbor searches in the first 30 principal components of the reference PCA space, capturing transcriptionally similar cells across datasets. These anchors were subsequently weighted according to their local neighborhood consistency and used to project query cells onto the reference UMAP embedding while preserving the structure of the reference manifold. Cell type labels were then transferred to query cells using a weighted aggregation of anchor-based label probabilities. All other parameters were set to their default values.

### RNA velocity

RNA velocity analysis was performed using scVelo (v0.2.5) on query datasets for which raw FASTQ files were available ^43^. Dataset SM_Yaman was excluded from the velocity analysis due to the absence of corresponding FASTQ files. For each included sample, velocyto (v0.17.17) was run on the Cell Ranger– derived BAM file, using the filtered barcode list and GENCODE GRCh38 GTF annotation to generate loom files containing spliced and unspliced transcript counts. Spliced and unspliced counts were extracted from the Cell Ranger–generated loom files for each time point and imported into scVelo. Genes were intersected across datasets to retain only shared features. To focus on developmental dynamics relevant to posterior axis elongation, the analysis was restricted to cell types shown in Fig. 2D, and further limited to cells assigned to axial-elongation–associated clusters based on label transfer from the human PCW3 reference. RNA velocity computation followed the standard scVelo workflow, including log-normalization and filtering of the data with default parameter, calculation of cell–cell relationships based on 30 principal components and identification of 30 nearest neighbors, stochastic modeling of transcriptional dynamics, and construction of the velocity graph. All other parameters were used at their default settings. Velocity stream plots were then visualized on the UMAP embeddings corresponding to axial-elongation–associated cells from PCW3, using the function of “scv.pl. velocity_embedding_stream”. For the RNA velocity vector projection to the UMAP of individual organoid in Fig. S5 batch correction for SM_Miao and TK_Rito was performed using the same methods and parameters as described in the original studies^27,29^. All other datasets were analyzed using the standard scVelo RNA velocity pipeline with 30 principal components and 30 nearest neighbors and the stochastic model, except for NT_Xue, for which the dynamical model was applied.

### Quantification of pathway-level signaling inputs

Small-molecule and growth factor treatments were converted into pathway-level signaling variables. For each signaling component, a raw input value was calculated as the product between factor concentration and exposure duration. Raw inputs were z-score scaled using the population standard deviation (ddof = 0). Pathway-level signaling variables were defined by summing the scaled inputs of components belonging to the same pathway (Fig. 4B). WNT was represented by normalized CHIR99021 input. FGF was defined as the sum of normalized FGF2 and FGF8 inputs. RA was defined as the sum of scaled RA and RAL inputs. SMAD inhibition was defined as the sum of scaled inputs from individual SMAD inhibitors (LDN193189, SB431542, DMH1, and A8301). SHH was represented by scaled SAG input. Five pathway-level variables were obtained for each protocol: WNT, FGF, RA, SMADi, and SHH. Pathway-level signaling variables were standardized prior to regression analysis using z-score scaling (StandardScaler).

### Quantitative Modeling of Signaling Pathways and Lineage Outputs

Principal component analysis (PCA) was used to explore variation in both lineage outputs and signaling variables across differentiation protocols. For PCA of tissue outputs, tissues were grouped into four categories: anterior neural tube (anterior NT, corresponding to forebrain–midbrain populations), posterior neural tube (posterior NT, encompassing hindbrain and cervical–thoracic populations), NMP, and mesoderm (including posterior PSM1, posterior PSM2, anterior PSM, somites, myocyte progenitors, mesoderm-derived muscle cells, and craniofacial mesoderm-like cells). Percentages of each tissues were standardized using z-score normalization prior to PCA, and the first two principal components were retained for visualization, enabling assessment of variation and similarities among protocols.

For PCA of signaling variables the z-scored scaled signaling pathway activity measures were used, with the first two principal components retained to visualize heterogeneity and relationships in signaling space across protocols.

### Multivariate linear regression analysis

Multivariate linear regression analyses were conducted in Python (v3.12.3), utilizing pandas for data management, numpy for numerical computations, scikit-learn for regression modeling, and matplotlib and seaborn for data visualization. Multivariate linear regression was performed using sklearn. linear_model.LinearRegression with the intercept included and default ordinary least squares settings, without regularization or cross-validation. For each lineage category (Anterior NT, Posterior NT, NMP, and Mesoderm), an independent regression model was fitted according to the following equation below:

**y=β0+β1WNT+β2FGF+β3RA+β4SHH**

Model performance was evaluated on the training data using the coefficient of determination (R²) and mean absolute error (MAE) as implemented in sklearn. metrics.

### Linear Regression Analysis

The analyses were conducted in Python (v3.12.3). For each signaling pathway and lineage percentage, univariate linear regression was performed to assess their relationship. Normalized signaling values and lineage percentages were used as the dependent variable. Linear regression lines were fitted using scikit-learn’s LinearRegression.

## Contribution statement

Y.W. conceptualization, methodology, investigation, data curation, analysis, visualization, and writing; R.B. conceptualization and methodology; C.H. data analysis; M.D. supervision and writing; C.S. supervision and writing.

## Acknowledgements

This work was performed using computational resources provided by the Academic Leiden Interdisciplinary Cluster Environment (ALICE) at Leiden University. This work was supported by the Lingling Wiyadharma Fonds ter bevordering van de beoefening der Natuurwetenschappen. W. Wang was supported by a fellowship from the China Scholarship Council (No. 202108210097).

## Notes

### Competing Interest Statement

The authors have declared no competing interest.

